# Reaching to sounds in virtual reality: A multisensory-motor approach to re-learn sound localisation

**DOI:** 10.1101/2020.03.23.003533

**Authors:** Chiara Valzolgher, Grègoire Verdelet, Romeo Salemme, Luigi Lombardi, Valerie Gaveau, Alessandro Farné, Francesco Pavani

## Abstract

When localising sounds in space the brain relies on internal models that specify the correspondence between the auditory input reaching the ears and initial head-position with coordinates in external space. These models can be updated throughout life, setting the basis for re-learning spatial hearing abilities in adulthood. This is particularly important for individuals who experience long-term auditory alterations (e.g., hearing loss, hearing aids, cochlear implants) as well as individuals who have to adapt to novel auditory cues when listening in virtual auditory environments. Until now, several methodological constraints have limited our understanding of the mechanisms involved in spatial hearing re-learning. In particular, the potential role of active listening and head-movements have remained largely overlooked. Here, we overcome these limitations by using a novel methodology, based on virtual reality and real-time kinematic tracking, to study the role of active multisensory-motor interactions with sounds in the updating of sound-space correspondences. Participants were immersed in a virtual reality scenario showing 17 speakers at ear-level. From each visible speaker a free-field real sound could be generated. Two separate groups of participants localised the sound source either by reaching or naming the perceived sound source, under binaural or monaural listening. Participants were free to move their head during the task and received audio-visual feedback on their performance. Results showed that both groups compensated rapidly for the short-term auditory alteration caused by monaural listening, improving sound localisation performance across trials. Crucially, compared to naming, reaching the sounds induced faster and larger sound localisation improvements. Furthermore, more accurate sound localisation was accompanied by progressively wider head-movements. These two measures were significantly correlated selectively for the Reaching group. In conclusion, reaching to sounds in an immersive visual VR context proved most effective for updating altered spatial hearing. Head movements played an important role in this fast updating, pointing to the importance of active listening when implementing training protocols for improving spatial hearing.

**HIGHLIGHTS:**

- We studied spatial hearing re-learning using virtual reality and kinematic tracking
- Audio-visual feedback combined with active listening improved monaural sound localisation
- Reaching to sounds improved performance more than naming sounds
- Monaural listening triggered compensatory head-movement behaviour
- Head-movement behaviour correlated with re-learning only when reaching to sounds

## 1. INTRODUCTION

Spatial hearing is a remarkable ability of the brain. To determine the spatial coordinates of sounds in the environment, the cognitive system exploits auditory cues resulting from the interactions between sound waves, the head and outer ears (Middlebrooks, 2015; Middlebrooks & Green, 1991; Wallach, 1940). Weighted combination of these auditory cues allows localisation of auditory events in the horizontal and vertical planes, as well as discrimination of their distance. Auditory cues change during the life span, due anatomical modifications of one’s own body during development (Clifton, Gwiazda, Bauer, Clarkson & Held, 1998) or changes in hearing threshold with ageing (Cranford, Andres, Piatz & Reissig, 1993; Dobreva, O’Neill & Paige, 2010). In addition, they change on a daily basis, due to the diversity of listening contexts to which we are all exposed (Majdak, Goupell & Laback, 2010). To cope with these continuous changes in auditory cues, the brain remains capable of updating sound-space correspondences throughout life (e.g., Carlile, Balachandar & Kelly 2014; Keating & King, 2015; Rabini, Altobelli & Pavani, 2019; Strelnikov, Rosito & Barone, 2011; Van Wanrooij, John & Opstal, 2005). In the present work, we examined reaching to sounds in virtual reality as a multisensory-motor strategy for re-learning sound localisation in adulthood.

Updating of sound-space correspondences typically occurs in a multisensory environment. Whenever the other sensory systems (e.g., vision) provide reliable spatial information about auditory events, the brain exploits these additional sensory sources to calibrate and optimize internal models for spatial hearing (Keating & King, 2015). This notion emerged, for instance, from studies that examined re-learning of sound-space correspondences when auditory cues were temporarily altered using monaural ear-plugs (Rabini et al., 2019; Strelninkov et al., 2011; Trapeau & Schönwiesner, 2015), ear molds (Van Wanrooij & Van Opstal, 2005) or non-individualised HRTFs (Head-Related Transfer Functions; Honda, Shibata, Gyoba, Saitou, Iwaya & Suzuki, 2007; Parseihian & Katz, 2012; Steadman, Kim, Lestang, Goodman & Picinali, 2019). In these simulated altered-listening conditions, multisensory training procedures proved effective for re-learning sound-space correspondences (for reviews see: Carlile, 2014; Keating & King, 2015; Knudsen & Knudsen, 1985; Mendonça, 2014; Irving & Moore, 2011). For instance, Strelnikov and colleagues (2011) studied the effects of audio-visual vs. auditory-only training on sound localisation in monaurally-plugged adults. They found that performance improvements were larger after a training that exploited spatially and temporally congruent audio-visual inputs, compared to a training based on auditory information alone. These findings complement other research in which participants received feedback after their sound localisation response, through audio-visual (Shinn-Cunningham, Durlach, & Held, 1998; Zahorik, Bangayan, Sundareswaran, Wang & Tam, 2006; see also Majdak, Walder & Laback, 2013) or visual cues (Bauer, Matuzsa, Blackmer & Glucksberg, 1966; Kumpik, Kacelnik & King, 2010; Mendonça, Campos, Dias & Santos, 2013). Finally, converging evidence in favour of multisensory-based training emerged from research in ferrets with unilateral and bilateral cochlear implants. Performance in early-deafened ferrets with bilateral cochlear implants improved consistently after a multisensory training that used interleaved auditory and visual stimuli (Isaiah, Vongpaisal, King, & Hartley, 2014).

In natural, off-laboratory conditions, spatial hearing is also an active task. Listeners spontaneously orient their head, eyes and trunk to the auditory source (e.g., when turning to listen to a speaker behind us), or they move their body in the environment to approach the auditory source (e.g., when reaching to grasp the mobile phone ringing on the table). These are all examples of “active listening” – a way of listening in which people are free to move head and body to interact with sound sources – which constitutes a distinguishing feature of all everyday listening conditions. Experimental evidence in support of the notion that active listening can improve sound localisation primarily comes from the study of head-movements during listening (Noble, 1981; Thurlow and Runge, 1967). Head-orienting to sounds is a spontaneous behaviour: it is detected in infants 2-4 days after birth (Muir & Field, 1979) and it follows the shortest path to the sound already between 6 and 9 months of age (van der Meer, Ramstad, & van der Weel, 2008). Adults have different propensities for head-movement (Fuller, 1992), but they orient their head to sounds easily when required by the task (Brimijoin, McShefferty, & Akeroyd, 2010). Most relevant for the present study, there is evidence that head-movement strategies change in altered hearing conditions. For instance, Brimijoin and colleagues (2010) showed that people with hearing-impairment have more complex head-movements. Also, people with asymmetrical hearing impairment use head-orienting to maximize the level of a target sentence in their better ear to solve a speech in noise task (Brimijoin & Akeroyd, 2012). Yet, studies that examined spatial hearing re-learning have typically limited the possibilities for head-movements and, in general, the contribution of active listening. Head-movements have been prevented by using a chinrest (Kumpik et al., 2010; Strelnikov et al., 2011) or by instructing participants to refrain from head-movements during listening (Rabini et al., 2019; Van Wanrooij & Van Opstal, 2005; Zahorik et al., 2006).

Further examples of active listening can be found in the few studies that investigated the role of gamified scenarios and reaching to sounds in acoustic space re-learning. Researches have asked participants to move a hand-held tool to hit a sound presented in virtual auditory space, while the head and the body are free to move (Honda et al., 2007; Honda et al., 2013; Ohunchi, Iwaya, Suzuki & Minekata, 2005). For instance, Parseihian and Katz (2012) explored the effect of a multi-modal training platform in which participants were involved in an active game-like scenario. They actively searched for animal sounds, scanning the space around them (front and back), through a hand-held track-ball. Again, participants were free to move their head. This training improved vertical localisation performances and reduced front/back confusion errors. A more recent study (Steadman et al., 2019) combined gamification, virtual reality and active listening (specifically, free head-movements) in a training protocol aimed at ameliorant sound localisation with virtual sounds. These authors trained participants to adapt to non-individualized HRTFs using virtual reality. They asked participants to listen to stationary virtual sounds either with the head static or free to move. Compared to the static listening condition, active listening improved sound localisation even over a very short timescale, corroborating the importance of head movements when re-learning sound-space correspondences.

These studies provide initial evidence in favour of active listening during re-learning of sound-space correspondences. Yet, they have not examined the specificity of reaching-to-sounds in these improvements, using comparable control conditions in which the listening context is kept identical while the reaching component is removed. Furthermore, despite they all introduced free head-movements during sound presentation and tracked them in real-time, they did not investigate the qualitative and quantitative nature of these head movements (e.g., head-movement number or amplitude, head-movement strategies). As such, these studies provide limited evidence of the actual role of head-movements in re-learning sound localisation.

Here we aimed to fill this gap by testing directly the hypothesis that reaching to sounds can promote updating of correspondences between auditory cues and external space more than a less active listening condition. To this aim, we asked two groups of participants to perform a sound localisation task in an interactive virtual reality scenario, while listening to real free-field sounds with one ear plugged. This simulated monaural listening condition requires participants to adapt to novel auditory cues while localising sounds in space. Participants wore a Head-Mounted Display (HMD) and perceived themselves in the centre of a virtual room. A semicircle of 17 visible speakers, each labelled with a different number, appeared in front of them within reaching distance in virtual reality. In each trial, a sound emitted from a real speaker appeared to originate from one of the visible speakers – yet its position could only be identified through auditory cues. The experimental group was instructed to stop sound emission by reaching to the correct source using a hand-held controller (reaching group). If the participant reached to the correct speaker the sound stopped, promoting a sense of agency over the auditory change. If instead the participant reached towards the wrong speaker the sound continued playing and the correct speaker started flashing, providing multisensory cues to actual sound location until the participants gave the correct response. The control group received identical stimulation and identical feedback, but was instructed to stop sound emission by naming the number that identified the active speaker (naming group). If active listening promotes re-learning of sound-space correspondences, performance in the monaural listening condition should improve faster for the reaching compared to the Naming group.

As additional aim, we examined the quantitative and qualitative nature of head-rotations as a function of response instructions (reaching/naming) and studied to what extent changes in head-movement behaviour related with sound localisation performance. Although head-movements were unconstrained for both groups, we hypothesised that head-movements could be facilitated in the Reaching group compared to the Naming group. The main rationale for this prediction stems from the literature on reaching and pointing to visual targets, which has documented a tight link between reaching and pointing actions and coordinated movements of the head and eyes (Biguer, Prablanc & Jeannerod, 1984; Fogt, Uhlig, Thach & Liu, 2002; Vercher, Magenes, Prablanc & Gauthier, 1994). Finally, we examined the impact on participants of our novel virtual reality scenarios and to what extent the two response instructions led to different engagement with the task.

## 2. MATERIAL AND METHODS

### 2.1. Participants

Twenty-eight participants (age: M = 20.93, SD = 2.48, range [18-30], 5 males, 14 right-handed in the Reaching Group, 13 right-handed in the Naming Group) were recruited to participate in the experiment, mostly among undergraduate students at the University of Trento. All participants signed an informed consent before starting the experiment, which was conducted according to the criteria of the Declaration of Helsinki (1964, amended in 2013) and approved by the ethical committee at the University of Trento (protocol: 2018-018). All had normal or corrected-to-normal vision, and reported no movement deficit. Hearing threshold was measured using an audiometer (Grason Stadler GSI 17 Audiometer) for all participants, testing different frequencies (250, 500, 1000, 2000, 4000 Hz), on the right and left ear separately. All participants had an average threshold below 11.7 dB HL.

### 2.2. Apparatus and stimuli

Virtual reality (VR) and kinematic tracking was implemented using an HTC Vive System, comprising one head-mounted display (HMD, resolution: 1080 × 1200 px, Field Of View (FOV): 110°, Refresh rate : 90 Hz), 1 controller (which was used by participants to interact with sounds), 1 tracker (which was placed above the speaker to track its position in real time) and 2 lighthouse base stations (which served for regular scanning of the position of the controller and the tracker). Tracking precision and accuracy of the HTC Vive System, as recently measured by our research group (Verdelet et al., in press), is adequate for behavioural research purposes. Specifically, the HTC Vive has submillimiter precision (0.237 mm) and near centimetre accuracy (9.0 mm when trackers are static; 9.4 mm when trackers are dynamic). The HMD was equipped with an SMI eye-tracking system (250 Hz). All stimuli were controlled and delivered using a LDLC ZALMAN PC (OS: Windows 10 (10.0.0) 64bit; Graphic card: NVIDIA GeForce GTX 1060 6GB; Processor: Intel Core i7-7700K, Quad-Core 4.2 GHz/4.5 GHz Turbo - Cache 8 Mo - TDP 95W) using Steam VR software and the development platform Unity.

The experiment was entirely run in a 4×3 meters ordinary room almost devoid of furniture, not treated for being anechoic and quiet. The subject sat in the centre of the room, with no constraints for her head (no chin-rest was used). The virtual environment in which participants were immersed was an ordinary squared room (grey wall), similar to the real experimental room. The room was empty and there were not objects on the walls, except a door which was placed in the back wall. In front of participants 17 visible speakers were positioned at ear level. In each session, subjects wearing the HDM saw themselves in the virtual scenario, sitting 50 cm from the centre of a semicircle of 17 speakers, spanning about ±80° of visual angle. Auditory stimuli were delivered by a real, unseen speaker (JBL GO Black), whose position was tracked in space. The auditory stimulus was an amplitude-modulated (4 Hz) white noise bursts (about 60 dB SPL as measured by a Decibel Meter TES1350A placed at ears level).

This solution allowed us the possibility of tracking the position of the tracker (speaker), the controller (handled by the participant) and the Head Mounted Display, via sensors on the headset and VR controllers using IR led positioned on the base stations (frequency sample 250 Hz). We design the software to allow the experimenter to track and align the real loudspeaker (the sound source) with the desired position in the virtual environment (i.e., in correspondence of one of the 17 loudspeakers).

To simulate monaural hearing experience, the right ear of each participant was occluded using an ear-plug (3M PP 01 002; attenuation value for high frequencies = 30 dB SPL; attenuation value for medium frequencies = 24 dB SPL; attenuation value for low frequencies = 22 dB SPL; Single Number Rating = 32 dB SPL; as reported by the manufacturer) and a unilateral ear muff (3M 1445 modified to cover only the right ear; attenuation value for high frequencies = 32 dB SPL; attenuation value for medium frequencies = 29 dB SPL; attenuation value for low frequencies = 23 dB SPL; Single Number Rating = 28 dB SPL; as reported by the manufacturer).

### 2.3. Procedure

We measured auditory performance during five different blocks, under two different listening conditions: binaural listening (B) and simulated monaural listening (M) in which the left ear was plugged (Figure 1A). In each block, participants performed 51 trials (3 trials for each source) of a sound localization task. First, participants were invited to sit down on a chair and wear the Head Mounted Display (HMD) and to perform an eye-tracker calibration (they had to follow a white dot moving on the screen with eyes). Then, they were immersed in the virtual scenario. Around them, the cylindrical speakers were located at the height of their ears. Participants were instructed that the starting position consisted in gazing at the central speaker (marked with a cross) and refrain from moving their head until the beginning of the sound. The system delivered the sound only when head (HMD) and eyes (tracked) were directed toward the centre. In such a way, we obtained replicable head and eyes posture across trials and participants. As soon as the sound started, participants were free to move their eyes and head as they wanted.

**Figure 1.**
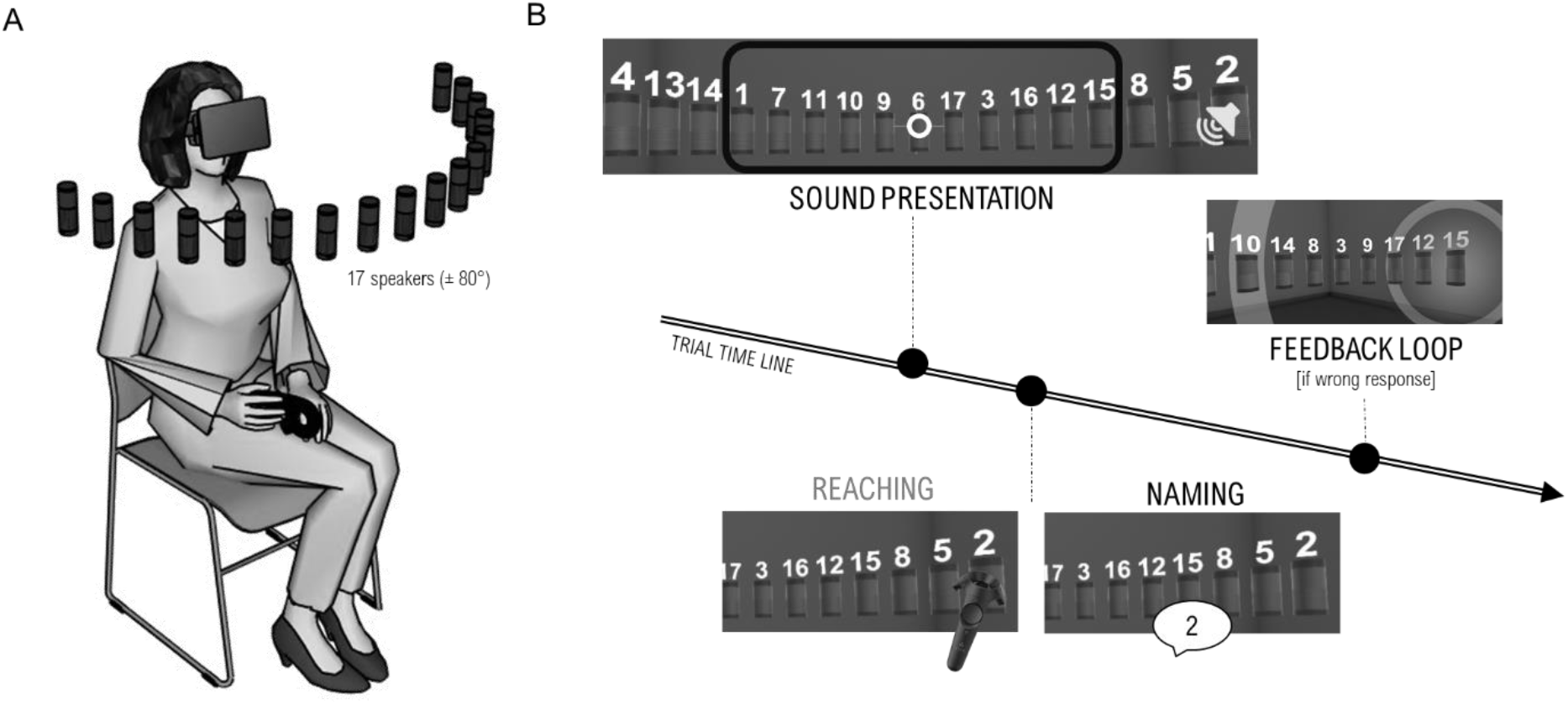
Experimental procedure and setting. (A) Schematic representation of participant and speaker positions. (B) Schematic representation of 1 trial: participants saw the virtual room and part of the speakers’ array (the black square represents participants starting view), they can turn head and move eyes to see all speakers and they stopped the sound moving the controller toward the source (Reaching group) or naming the label (Naming group). If their answer was wrong, the correct source started to flash providing a feedback.

After each block, participants were given breaks and were asked to judge their performance (“How do you judge your performance?” from 1 = really bad to 9 = really good) and their perceived effort during the block (“How much effort (in terms of energy and cognitive resources) did the task require?” from 1 = none to 9 = many). At the end of 5 blocks, participants were invited to complete a questionnaire presented using Google Forms. The questionnaire comprised general questions about personal data (i.e., age, gender), questions to investigate their relationship with sound during life, specific questions to deepen the quality of our VR technology and its ergonomics, a question concerning their strategy to perform the task, two questions concerning the feeling of being able to switch the sound off (Agency) and 15 items of 2 scales (Focus of Attention and Satisfaction), used typically to measure the engagement in video game-based environments (Figure 2). We adapted these items from the work of Wiebe and colleagues (2013) to make them more suitable for our VR experience. Participants indicated their agreement with each item using a 7-points Likert scale (from 1 = totally disagree to 5 = totally agree).

**Figure 2.**
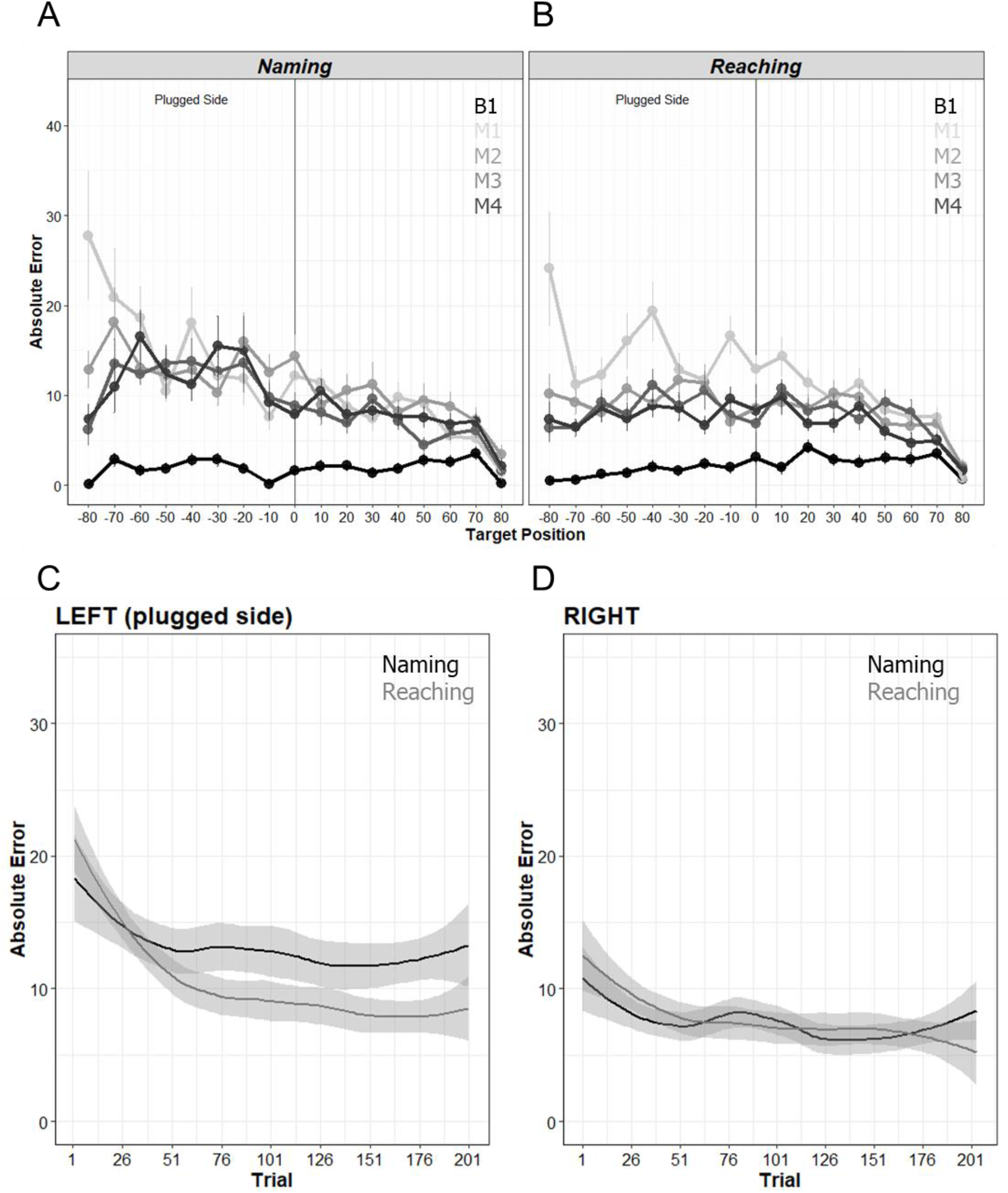
A and B) Absolute errors (in degrees) as a function of Blocks (B = binaural block, M1 = first monaural block, M2 = second monaural block, M3 = third monaural block, M4 = fourth monaural block) and of group (Naming (A) and Reaching (B)). Graphs bars represent standard errors. C and D) Absolute errors as a function of trial for Naming (black) and Reaching (grey) group when the target sound came from the left (plugged side) (C) and right (D) side of the space.

We divided participants in 2 groups: the Reaching group and the Naming group (Figure 1B). The Reaching group performed the task by moving a controller with their dominant hand to touch the speaker emitting the sound. The Naming group performed the same task by reading aloud the number located above the presumed source speaker (see Fig 1B). If the response was correct, the sound immediately stopped. If the response was wrong, the correct speaker produced a pulsing red light while the sound continued. This provided participants with audio-visual feedback about the correct position of the sound source. Participants either moved the controller in contact with the correct source or named the number of the right speaker. When the correct speaker was reached (or named) the sound stopped. Importantly, participants were informed that they were free to exploit head-movements while listening to target sounds, without time pressure on their response. Both reaching and naming responses were followed by a brief vibration of the hand-held controller (which was kept in hand also during the naming task).

#### Head-movement pre-processing

To study head-movements, we calculated the tangential velocity on the x, y, z axis (expressed in degrees of rotation) using two-points central difference derivate algorithm (Bahill & McDonald, 1983) with 5 points for the half-window. The onset and the end of the movements were computed automatically using a velocity-based threshold (10°/s). Head-movements were checked manually by visualizing the spatial rotation changes of the head and its speed using a home-made tool box in MATLAB R2018b. Subsequently, we applied a duration-based filter (> 350 millisecond) and a spatial-based filter (> 10°).

## 3. RESULTS

### 3.1. Sound localisation performance

#### 3.1.1. Ear-plug effect

To describe the immediate effects of monaural plugging, we compared absolute errors before and after ear-plugging. Absolute errors (calculated in each trial as the difference in degrees between actual and reported azimuth position of the sound) were entered into an Analysis of Variance (ANOVA) with TARGET POSITION (17 positions, from −80° to +80°) and LISTENING CONDITION (binaural B, monaural M1) as within-participants variables and GROUP (reaching, naming) as between-participants variable. All statistical analyses using the software JASP 0.9.1.0 and R-studio (version 1.0.143).

The analysis revealed a main effect of LISTENING CONDITION, *F*(1,26) = 64.79, *p* < .001, *η*^*2*^ = .71, a main effect of TARGET POSITION, *F* (3.95, 102.60) = 5.96, *p* <.001, *η*^*2*^ = .18, and the 2-way interaction between LISTENING CONDITION and TARGET POSITION, *F* (3.9, 101.51) = 7.36, *p* < .001, *η*^2^ = .22. Sound localisation error increased significantly in M1 compared to the B listening condition (see black and light grey lines in Figures 2A and 2B), particularly for target delivered more towards the plugged ear (left). This effect emerged irrespective of GROUP (main effect and all interactions, p-values > .27).

Participants were aware of their performance change after monaural plugging. Performance assessment scores (“*How do you judge your performance?*” from 1 = really bad to 9 = really good) were lower in M1 (Reaching: M = 3.29; DS = 1.90; Naming: M = 3.50, DS = 1.40) compared to the B listening condition (Reaching: M = 6.86; DS = 1.35; Naming: M = 6.57, DS = 0.85; Chi-square = 24.14, *p* < .001, in non-parametric Friedman test), irrespective of GROUP. Participants also reported that monaural plugging required additional cognitive effort. Effort assessment scores (“*How much effort (in terms of energy and cognitive resourced used) did the task require?*” from 1 = none to 9 = many) were larger in M1 (Reaching: M = 6.71; DS = 1.33; Naming: M = 6.21, DS = 1.72) compared to the B listening condition (Reaching: M = 3.29; DS = 1.49; Naming: M = 3.57, DS = 1.95; Chi-square = 28, *p* < .001, in non-parametric Friedman test), again irrespective of GROUP.

#### 3.1.2. Re-learning

To study re-learning across successive monaural blocks, we entered absolute errors into an ANOVA with TARGET POSITION (17 positions, from −80° to +80°) and MONAURAL BLOCKS NUMBER (M1, M2, M3 and M4) as within-participants variables, and GROUP (reaching, naming) as between-participants variable. This analysis revealed a main effect of MONAURAL BLOCK NUMBER, *F*(3,78) = 8.51, *p* < .001, *η*^2^ = .24, a main effect of TARGET POSITION, *F* (16, 416) = 8.89, *p* <.001, *η*^2^ = .24, and the 2-way interaction between MONAURAL BLOCK NUMBER and TARGET POSITION, *F* (48, 1248) = 3.02, *p* < .001, *η*^2^ = .10. Participants improved across M blocks, especially for target positions more towards the plugged ear (left) (Figures 2A and 2B). No main effect or interaction involving the GROUP variable emerged (all p-values > .18), when considering overall performance in a block.

To further investigate changes in localisation error as a function of practice in the two groups, we analysed performance changes across the 204 trials (51 trials in each of the 4 blocks) using a linear mixed model (LMM). Specifically, we entered in the model the fixed effects and the interactions of three components: GROUP (naming, reaching), PRACTICE (as a continuous variable of 204 items) and STIMULATION SIDE (left, right; note that trials from the central speaker were excluded from this analysis). Participant and target position were considered as random effects in the model. This analysis revealed that absolute errors were influenced by STIMULATION SIDE (*t* = −6.235, *p* < .001): as expected, errors were more pronounced on the side ipsilateral to the plug (left; see Figures 3C and D). Sound localisation performance was also influenced by PRACTICE (*t* = −4.643, *p* < .001), revealing that participants learned across monaural trials. Crucially, our model showed also learning across trials was influenced by the type of task performed by participants (i.e., GROUP; *t* = −2.861, *p* < .01). The Reaching group learned more rapidly and reduced localisation errors to a greater extent compared to the Naming group. Error reduction was more pronounced for sounds ipsilateral to the plug (left) compared to contralateral ones (*t* = 2.521, *p* < .05).

**Figure 3.**
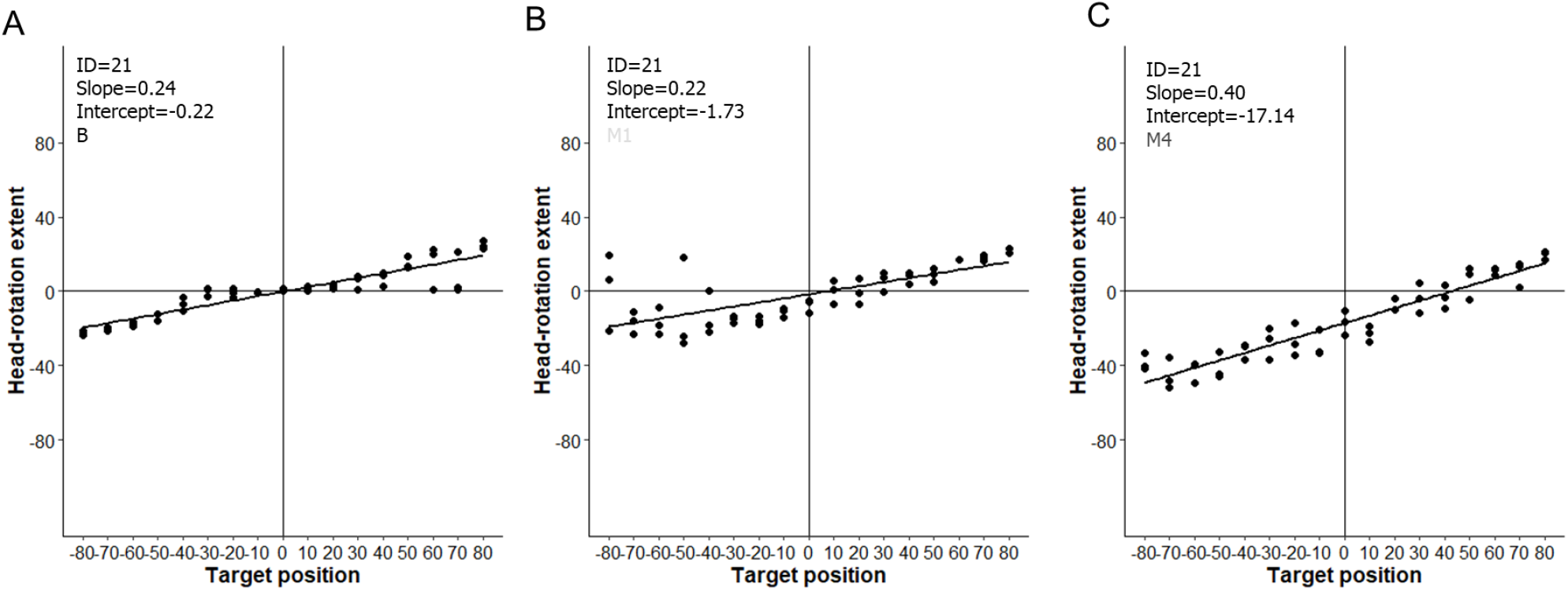
Head-rotation extent as a function of target position of participants 21 during first binaural block B (A), the first monaural block M1 (B) and the last monaural block M4 (C).

Participants were aware that their performance improved across monaural blocks. Performance self-assessment scores increased across monaural blocks (M4: Reaching: M = 4.29; DS = 1.38; Naming: M = 3.93, DS = 1.69; Chi-square = 10.02, *p* < .001, in non-parametric Friedman test), irrespective of GROUP. Participants did not report any change in terms of energy and cognitive resources used to do the task across monaural blocks: effort assessment scores did not change across monaural blocks in either GROUP (all *ps* > .23).

Finally, following previous works (Rabini et al., 2019; Steadman et al., 2019; Valzolgher, Campus, Rabini, Gori & Pavani, under review) we examined whether re-learning across blocks relates to the magnitude of the immediate ear-plug effect. To this aim, we correlated across participants the ear-plug effect (measured as the difference in absolute localisation error between B and M1; positive values indicate larger plug-effect) and re-learning (measured as the difference in absolute localisation error between M1 and M4; positive values indicate larger re-learning). Significant correlations were observed (*r* = 0.55, *p* = 0.003), in agreement with the notion that larger re-learning emerges in those participants who experienced the larger plug-effect.

### 3.2. Head movements

To study head-movements we considered 5 dependent variables, obtained either from single trial measures or across all trials in a block. Single trial measures were: (1) number of head-movements per trial, (2) head-movement amplitude per trial (in degrees), and (3) average head-movement duration per trial (in seconds). Measures across trials were: (4) head-rotation around vertical axis as a function of target position, and (5) overall head-direction bias during an entire block.

The latter two indices were extracted by linear fitting of the changes in head-rotation extent as a function target position (see example in Figure 3). Head-rotation extent was calculated as the average between the leftmost and rightmost head-rotation, for any given target position. The slope of the fitting line captures head-rotation around the vertical axis: the larger the slope the more participants rotated their head as a function of target eccentricity. In the representative participant (ID=21) shown in Figure 3, the slope remained stable from B to M1 (0.24 vs. 0.22) but increased from M1 to M4 (0.22 vs. 0.40), indicating a larger propensions to rotate the head towards peripheral targets in M4 compared to M1. The intercept of the fitting line captures the overall head-direction bias during an entire block, with negative values indicating a bias towards the left (plugged) side. In the same representative participant, the intercept became progressively more negative from B to M1 to M4 (−0.22, −1.73, −17.14, respectively) revealing the bias of this participant to rotate the head further to the plugged side, thus bringing the unplugged ear towards the target.

#### 3.2.1. Ear-plug effect

To describe the immediate effects of monaural plugging on head-movements, we compared the number of head-movements per trial before and after ear-plugging. The number of movements (calculated in each trial as described in pre-processing and analyses paragraph) was entered into an ANOVA with TARGET POSITION (17 positions, from −80° to +80°) and LISTENING CONDITION (B, M1) as within-participants variables and GROUP (reaching, naming) as between-participants variable. The analysis revealed a main effect of LISTENING CONDITION, *F*(1,26) = 48.65, *p* < .001, *η*^2^ = .63, caused by more head-movements in M1 compared to the B listening condition (see black and light grey lines in Figures 4A and B). A main effect of TARGET POSITION, *F* (8.61, 223.82) = 10.30, *p* <.001, *η*^2^ = .28, reflecting more head-movements for peripheral targets overall. Finally, there was a 2-way interaction between LISTENING CONDITION and TARGET POSITION, *F* (16,416) = 5.51, *p* < .001, *η*^2^ = .17) caused by more head-movements in M1 compared to B especially when targets were more central (F (16,416) = 5.51, < .001, η^2^ = .17). This effect emerged irrespective of groups (main effect and all interactions, p-values > .17).

**Figure 4.**
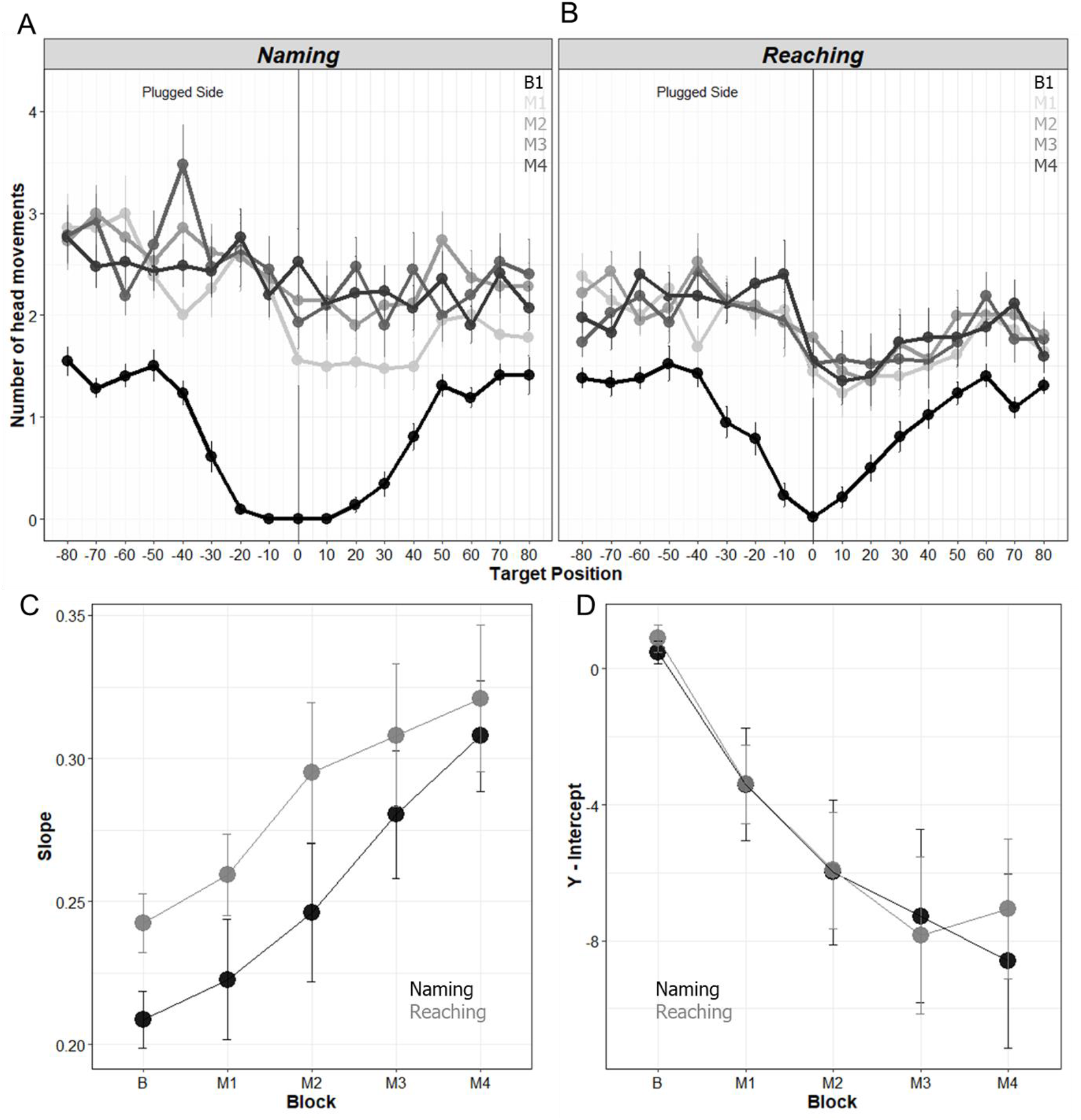
A and B) Number of movements as a function of Blocks (B = binaural block, M1 = first monaural block, M2 = second monaural block, M3 = third monaural block, M4 = fourth monaural block) and of group (Naming, A and Reaching, B). Graphs bars represent standard errors. C and D) Slope (C) and Intercept (D) values as a function of blocks B1 M1 M2 M3 M4), Naming (black) and Reaching (grey).

Similar analyses on head-movement amplitude per trial and average duration per trial revealed a main effect of LISTENING CONDITION (amplitude: *F*(1,26) = 73.05, *p* < .001, *η*^*2*^ = .73; duration: *F*(1,26) = 156.56, *p* < .001, *η*^*2*^ = .85), a main effect of TARGET POSITION (amplitude: *F* (7.42, 192.79) = 121.85, *p* <.001, *η*^*2*^ = .81, duration: *F* (7.07, 182.18) = 69.05, *p* <.001, *η*^*2*^ = .72) and an interaction between LISTENING CONDITION and TARGET POSITION (amplitude: *F* (5.70, 148.29) = 26.90, *p* < .001, *η*^2^ = .50, duration: *F* (6.19, 160.90) = 20.26, *p* < .001, *η*^*2*^ = .43). During M1, head-movement amplitude was larger and average head-movement duration was longer compared to the B listening condition, especially for targets more towards the plugged ear (left). Furthermore, we observed an interaction between TARGET POSITION and GROUP (F (7.42, 192.79) = 2.44, p = .018, η^2^ = .02), which revealed a general tendency of Reaching group to make wider head-movements when the sound came from the periphery.

To study how participants explored auditory space with their head during an entire block, we focused on (1) head-rotation around vertical axis as a function of target position (slope of the fitting line, as specified above) and (2) overall head-direction bias during an entire block (intercept of the fitting line). Both measures were entered separately into an ANOVAs with LISTENING CONDITION (B, M1) as within-participants variable and GROUP (reaching, naming) as between-participants variable. The analyses on head-rotation around vertical axis revealed that participants in the Reaching group turned their head as a function of target position to a greater extent compared to those in the Naming group (main effect of group, *F*(1,26) = 4.48, *p* = .04, *η*^*2*^ = .15). In other words, they explored auditory space with their head to a greater extent (compare B and M1 in Figures 4C). No main effect or interaction involving LISTENING CONDITION emerged. The analysis on the overall head-direction bias revealed a main effect of LISTENING CONDITION, *F*(1,26) = 18.64, *p* = .04, *η*^*2*^ = .42. Intercepts became negative in M1 (Naming: M = −3.41, SD = 6.20; Reaching: M = −3.39, SD = 4.31) compared to the B listening condition (Naming: M = 0.50, SD = 1.26; Reaching: M = 0.89, SD = 1.48). This means that, when listening with the left ear plugged, all participants turned their head leftward, to approach the sound with their unplugged (right) ear (compare B and M1 in Figure 4D).

#### 3.2.2. Re-learning

To study the effect of re-learning on head-movements, we entered number of head-movements in an ANOVAs with TARGET POSITION (17 positions, from −80° to +80°) and MONAURAL BLOCKS NUMBER (M1, M2, M3 and M4) as within-participants variables and GROUP (reaching, naming) as between-participants variable. The analysis revealed a main effect of TARGET POSITION, *F* (16,416) = 11.49, *p* <.001, *η*^*2*^ = .14 and an interaction between MONAURAL BLOCKS NUMBER and TARGET POSITION, *F* (48, 1248) = 1.37, *p* = .048, *η*^*2*^ = .048). Sound delivered from positions closer to the plugged ear caused more head-movements compared sounds closer to the unplugged ear, selectively for blocks M1 and M2 (simple main effect: p<0.01 and p=0.02, respectively). This effect emerged irrespective of group (main effect and all interactions, p-values > .26). (Figures 4A and B).

Similar ANOVAs on head-movements amplitude and head-movement duration revealed a main effect of MONAURAL BLOCK NUMBER (amplitude: *F*(3,78) = 3.63, *p* = .02, *η*^*2*^ = .12; duration: *F*(3,78) = 8.47, *p* < .001, *η*^*2*^ = .24), a main effect of TARGET POSITION (amplitude: *F* (16, 416) = 39.31, *p* <.001, *η*^*2*^ = .59, duration: *F* (16, 416) = 26.72, *p* <.001, *η*^*2*^ = .50), but no interaction (all ps > .25). Participants increased head-movement amplitude and head-movement duration across the successive monaural blocks. As expected from the outcome of the earlier analyses, both indices were larger for sounds ipsilateral to the plug compared to contralateral ones.

Finally, we examined how participants explored auditory space with their head across monaural blocks. Slopes and intercepts across participants were entered into separate ANOVAs with MONAURAL BLOCK NUMBER (M1, M2, M3 and M4), as within-participants variable and GROUP (reaching, naming) as between-participants variable. Both analyses revealed only a main effect of MONAURAL BLOCK NUMBER (slope: *F*(3,78) = 12.56, *p* <.001, *η*^*2*^ = .32; intercept: *F*(1.63,42.44) = 10.71, *p* <.001, *η*^*2*^ = .29). As monaural blocks progressed, participants in both groups increased their exploration of auditory space (Figures 4C) and turned their head more to approach sounds with their unplugged (right) ear (Figure 4D).

### 3.3. Relation between changes in sound localisation and space explored with the head

#### 3.3.1. Ear-plug effect and space explored with the head

As detailed above, participants changed the portion of space explored with the head (as index by the slope value) when listening with one ear plugged (see section 3.2.1). We examined if the immediate ear-plug effect (i.e., changes in absolute localisation error between M1 and B; positive values indicate larger plug-effect) correlated with changes in the space explored with the head (i.e., changes in slope between M1 and B). This correlation is shown in Figure 5A, separately for each group (Pearson correlation with Bonferroni correction: reaching, r = 0.70, p = 0.01; naming, r = 0.79, p = 0.001). Irrespective of group, the more participants increased explored space between B and M1 blocks, the less the cost of ear plugging on their sound localisation performance.

**Figure 5.**
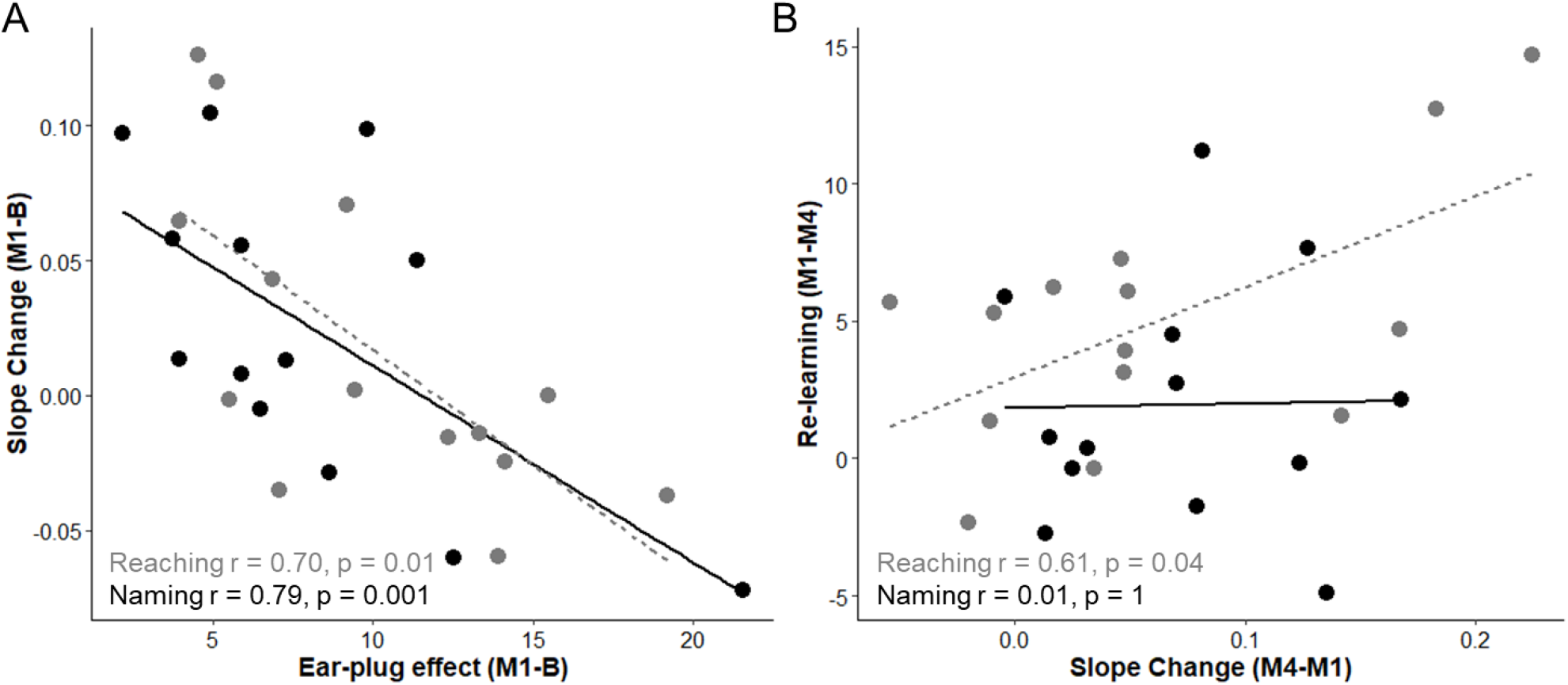
A) Correlation between Ear-plug effect and Slope Change. B) Correlation between Slope Change and Re-learning across monaural blocks. In both graphs participants in the Reaching group are indicated in grey (regression line shown as dashed) and those in the Naming group are indicated in black (regression line shown as continuous).

#### 3.3.2. Re-learning and space explored with the head

The portion of space explored with the head also changed across monaural blocks (comparison between M1-M4 blocks, see section 3.2.2). We examined if the immediate re-learning (i.e., changes in absolute localisation error between M4 and M1; positive values indicate larger improvement after trials repetition) correlated with changes in the space explored with the head (i.e., changes in slope between M4 and M1). This correlation is shown in Figure 5B, separately for each group (Pearson correlation with Bonferroni correction: reaching, r = 0.61, p = 0.01; naming, r = 0.01, p = 1). A positive relation emerged for the Reaching group, but not for the Naming group. Comparing correlations across group (as in Eld, Gollwitzer and Schmidt, 2011) a significant between-group difference emerged (z = −1.64, p = 0.05). This evidence showed that compensatory head-movement behaviour correlated with re-learning selectively for participants in the Reaching group.

### 3.4. VR experience

At the end of the experiment, participant replied to questions about their experience using VR system, to assess also the feasibility of our approach. They judged that the overall experience with virtual reality was positive (M = 7.86, SD = 1.15, using a scale from 1 =negative to 9 = positive), that the scene appeared realistic (M = 6.61, SD = 1.47, using a scale from 1 = unrealistic to 9 = very realistic) and reported no feeling of losing balance (M = 1.53, SD = 1.07, using a scale from 1 = not at all to 9 = completely) or being annoyed (M = 2.61, SD = 1.81, using a scale from 1 = not at all to 9 = completely).

Participants were also queried about the sense of agency and the perceived gaming experience when performing the two tasks (i.e., reaching or naming).

#### Sense of agency

Participants indicated their agreement with 2 items (“I felt that I can turn off the sound”, “I felt that I can act on the sound”) using a 5-points Likert scale (from 1 = totally disagree to 5 = totally agree). Participants in both groups felt equally that they could act on sounds (reaching: M = 2.29, SD = 0.99; naming: M = 1.71, SD = 0.99; *W* = 64, *p* = .14 on Mann Whitney). However, the Reaching group reported the experience of turn off the sound strongly (M = 4.43, SD = 0.51) compared to the Naming group (M = 1.21, SD = 0.58), *W* = 0.00, *p* < 0.001 (Mann Whitney).

#### Gaming experience

To investigate their gaming task-related experience, we proposed 15 items of 2 scale of the User Engagement Scale (UES) questionnaire, typically used to measure engagement during video-game playing (Wiebe, Lamb, Hardy & Sharek, 2014): focus of attention scale and satisfaction scale (Figure 6). We compared responses of the two groups for each item of the 2 scales, as well as the cumulative indices (average of the respective items, Cronbach’s α: 0.8; 0.89) in t-test analyses. Overall, our virtual reality scenario was an effective method to create a game-like situation, capable of actively involving participants. The satisfaction scale items revealed a difference between tasks: participants of the Reaching group (M = 3.71, SD = 0.83) perceived the experience more rewarding compared to the Naming group (M = 2.93, SD = 0.83; *W* = 47.50, *p* = .015). Moreover, they expressed more interest in doing the task (Reaching: M = 4.71, SD = 0.47; Naming: M = 4.07, SD = 0.92, *W* = 60, *p* = .05).

**Figure 6.**
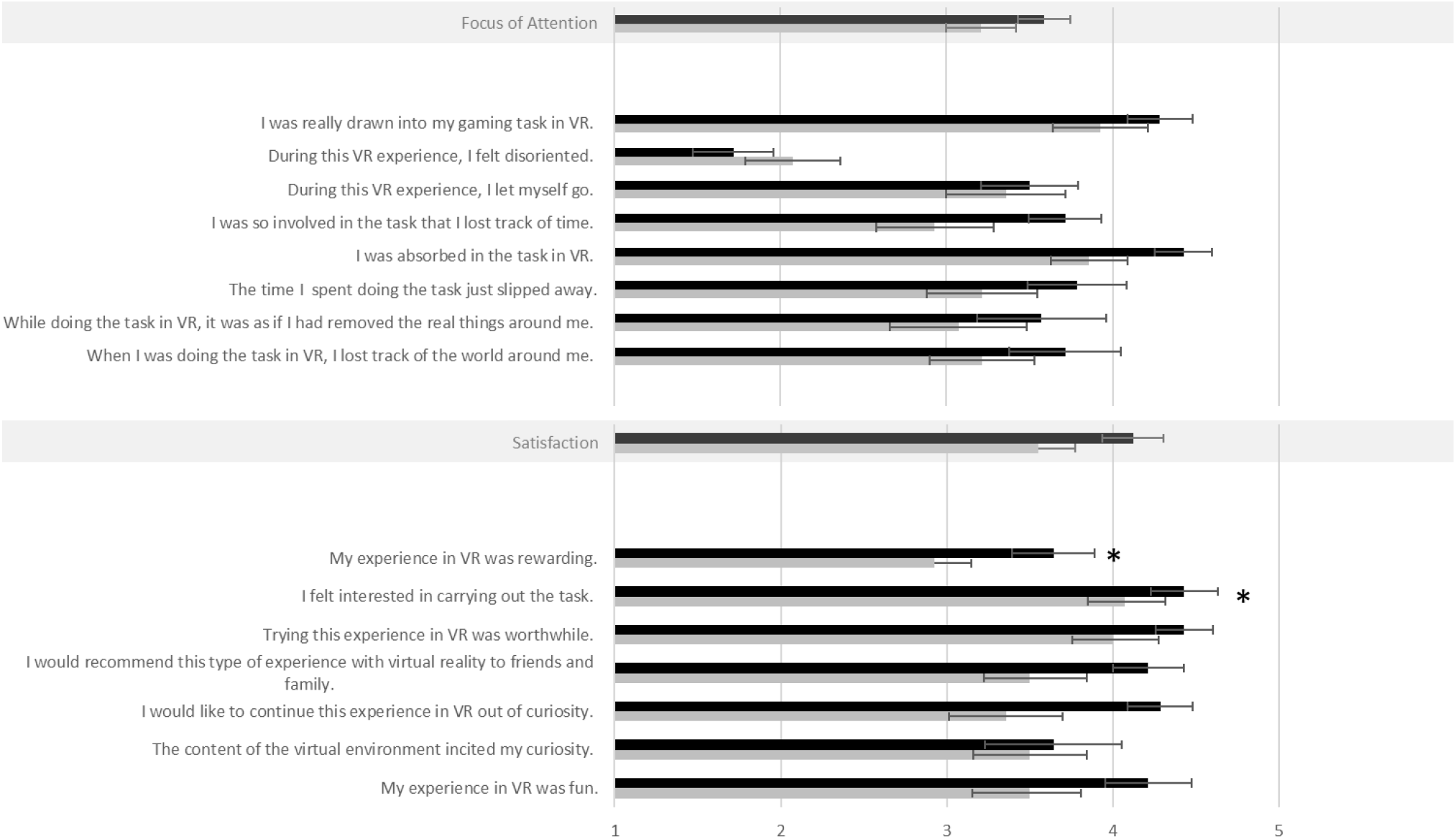
Rating scores on the experience with VR questionnaire, as a function of Reaching (black) or Naming (grey) group. Cumulative scores for focus of attention and satisfaction are reported highlighted in grey. Thicker bars represent mean values and smaller bars represents standard errors.

## 4. DISCUSSION

The ability to localise sounds relies on internal models that specify the correspondences between the auditory input reaching the ears and initial head-position with coordinates in external space. These correspondences can be updated throughout life (Carlile, 2014; Keating & King, 2015; Rabini et al., 2019; Strelnikov et al., 2011; Van Wanrooij et al., 2005), setting the basis for re-learning spatial hearing abilities in adulthood. This is particularly important for individuals who experience long-term auditory alterations (e.g., hearing loss, hearing aids, cochlear implants) as well as individuals who have to adapt to novel auditory cues when listening in virtual auditory environments. Until now, however, several methodological constraints have limited our understanding of the mechanisms involved in spatial hearing re-learning. Specifically, the potential role of active listening and head-movements have remained largely overlooked. Using a novel methodology, based on virtual reality and real-time kinematic tracking of the participant’s head and hand, we examined the contribution of active multisensory-motor interactions with sounds in the updating of sound-space correspondences.

Our experimental approach is based on a new apparatus (SPHERE; Pavani et al., 2017; Gaveau et al., in preparation) that allows to present free-field sounds from a real loudspeaker, placed at pre-determined locations and aligned with a virtual reality scenario. This allows full control over visual cues about the visual environment (here, an empty room), cues to sound position (here, a visible array of virtual loudspeakers) and feedback (here, the flashing speaker that appeared in case of mistaken responses). Most importantly, our approach allows full monitoring of head-position. This ensured identical straight-ahead initial posture for all trials and participants, but also permitted tracking of spontaneous head-movements during the response phase. Finally, precise kinematic of the hand response was measured in real-time. This allowed measuring of end-pointing positions in the reaching group and ensured compliance with the instructions in the Naming group.

Four novel findings emerged. First, we documented that updating of sound-space correspondences during monaural listening can occur over a relatively short period of time (about 200 trials, completed in approximately 50 minutes). Second, we showed that monaural plugging affected both sound localisation performance and spontaneous head-movements. Third, in line with our main prediction, we found that reaching to sounds proved more effective than naming sound position for updating altered spatial hearing. Fourth, we document that head-movements played an important role in this fast updating: reduction in localisation errors was accompanied by wider portion of space explored with the head. Notably, performance improvements and changes in head-movement extension correlated selectively for the reaching group, providing initial evidence for an interaction between reaching to sounds, head-movements and re-learning of sound-space correspondences.

### 4.1. Sound localisation improves rapidly during monaural listening

The present study demonstrates that normal hearing adults listening with one ear plugged and muffed can improve in sound localisation over a short period of time. Both group of participants (reaching and naming) progressively reduced their error over trial repetitions, within a single testing session. We favoured re-learning through a combination of audio-visual feedback (Strelnikov et al., 2011; Rabini et al., 2019) and active interactions with the sounds (spontaneous head-movements, agency over the sound), which made participants engaged and comfortable when performing the tasks. Despite being the first VR experience for most of the participants, the positive assessment they gave confirms its feasibility, which represents an important pre-requisite for the use of VR applications in auditory perception research and clinical rehabilitation (see 3.4).

While re-learning of sound-space correspondences has been previously documented (Carlile, 2014; Kacelnik, Nodal, Parsons & King, 2006; Keating & King, 2015; Nawaz, McNeill & Greenberg, 2014; Steadman et., 2019), to the best of our knowledge, fast improvement over trials have not been studied and described before. Traditionally, updating of space-sound correspondences has been examined in paradigms which comprised pre- and post-testing phases interspersed with training procedures which could occur over several days (e.g., 5 days in Strelnikov et al., 2011 and in Irving & Moore, 2011; 10 days in Ohuchi et al., 2005). Thus, even when acoustic space re-learning occurred after small number of short training session (3 sessions, lasting 15 minutes as in Steadman et al., 2019), they were performed across multiple days. Although in our study we did not test the generalisation of sound localisation re-learning, we investigated the processes of sound-space correspondences updating over the course of 4 monaural blocks of 51 trial each. We believe this fast change is more compatible with re-weighting of auditory cues, rather than actual re-mapping of sound-space coordinates. Yet, the present findings show that the processes of auditory cues re-weighting could occur in less than an hour (see Kacelnik et al., 2006; Irving and Moore, 2011; Kumpik et al., 2010).

One aspect worth mentioning is that the ear-plug effect (i.e., the extent to which performance changed from binaural to monaural listening) impacted on re-learning across blocks, indicating that larger re-learning emerges in those participants who experienced the larger plug-effect (for related findings see: Rabini et al., 2019; Steadman et al., 2019; Valzolgher et al., under review). As suggested by Rabini and colleagues (2019), the relation between ear-plug effect and re-learning may be the consequence of the perceived discrepancy between the response and the actual position of the sound. This discrepancy was continuously experienced during our tasks, whereby the visual feedback made actual sound position explicit. The more participants were aware of this discrepancy, the higher the possibility that a new correspondence between auditory cues and space could been stored. This finding converges in suggesting that feedback plays a critical role in acoustic space re-learning processes, even when experience is limited to few hundreds of trials (Rabini et al., 2019; Steadman et al., 2019; Valzolgher et al., under review).

### 4.2. Monaural listening triggered compensatory head-movements

The important role of head-movements in sound localisation has been emphasised since the 1940’s (Wallach, 1940), but studied sporadically until recently (Iwaya, Suzuki & Kimura, 2003; Morikawa & Hirahara, 2013; Perrett & Noble, 1997; Toyoda, Morikawa & Hirahara, 2011). To the best of our knowledge, this is the first study to examine changes in head-movement behaviour in monaural compared to binaural listening, and during re-learning of sound-space correspondences. Here, we characterised head-movements with 5 different variables (number of head-movements per trial, head-movement amplitude per trial, average head-movement duration per trial, head-rotation around vertical axis, and overall head-direction bias during an entire block) and showed that head-movements change as a function of listening condition as well as re-learning. Monaural plugging affected spontaneous head-movements, which increased and became wider over monoaural listening repetitions. We propose this was functional to deal with the altered listening condition and favoured the auditory cues re-weighting process.

Sound localisation is often accompanied by head orientation and movements that lead to localisation performance benefits in the azimuth and elevation planes (Morikawa & Hirahara, 2013; Perret & Noble, 1997; Thurlow & Runge, 1967) and facilitate disambiguating the front/back confusion (Brimijoin et al., 2012; Kim, Barnett-Cowan, & Macpherson, 2013; McAnally & Martin, 2014; Perret & Noble, 1997; Wightman & Kistler, 1999). In our study, participants moved their head to reach the most favourable listening position (as indexed by the slope, see 3.2.2). The need to cope with the altered acoustic space is fulfilled by increasing the frequency and extent of head-turns to favour perception with the unplugged ear. This strategy is comparable to the head-behaviour measured when hearing in noise: as reported by Brimijoin and colleagues (2012), when participants listen to sounds in noise, head-movements can be used to increase the level of the signal in their better ear, or to increase the difference in level between the signal and the noise. Remarkably, this type of head movement differs compared to those performed in more typical hearing situations (i.e., binaural hearing), whereby participants rather tend to face the speaker frontally when performing a sounds localisation task (Iwaya et al., 2003; Thurlow, Mangels & Runge, 1967).

### 4.3. Reaching to sounds promote re-learning more than naming

One key observation of our study is that reaching to sounds improved performance more than naming sounds. Participants who interacted with sounds by reaching them learned more rapidly and reduced localisation errors to a greater extent compared to those who interacted with sounds simply reading labels.

This observed advantage of reaching over naming is most likely the results of multiple factors. One first factor which has emerged from the present work, is the different role of head-movements in the two groups. Head-movement amplitude was overall wider for sounds at the periphery for reaching compared to naming participants, irrespective of listening condition (binaural vs. first monaural block; see 3.2.1). Most importantly, although participants in both groups were free to move the head during the task, a significant correlation between performance improvement and head-movements extension emerged selectively for the reaching group (see 3.3.2). Thus, reaching to sound seemed to be more effective in promoting active listening strategies involving head-movement, which in turn can impact on re-learning.

Other factors that may contribute to re-learning of sound-space correspondences when reaching to sounds should however be considered. First, reaching to sounds requires a coordination of different effectors (eyes, head, hand) into a common reference frame (Cohen & Andersen, 2002). In turn, this may result in a more stable (or salient) spatial coding of sound source location and favour re-learning of sound-space correspondences. Second, the documented feeling of agency on sound sources resulting from the gesture of reaching an object may have played a role, possibly leading to a stronger motivation when performing the task (Aarhus, Grönvall, Larsen, & Wollsen, 2011). In line with this speculation, note that the reaching group perceived the experience as more rewarding overall, and expressed more interest in the task compared the naming group. Third, reaching movement may have helped directing attention towards the position occupied by the sound source, because of additional visual, proprioceptive and kinaesthetic cues resulting from the action. Future works should explore the contribution of these additional factors in promoting re-learning of sound-space correspondences during active listening conditions involving reaching to sounds.

## 4.4. Conclusions

In this article we demonstrated the possibility to improve sound localisation abilities, even with limited experience. Most importantly, we documented the compensatory head-movement behaviour triggered by the monaural listening condition and we showed that reaching to sounds can promote re-learning of sound-space correspondences more than naming sounds. These findings highlight the benefits of reaching to sound as an active listening approach to re-learning sound-space correspondences.

The implications of these results are two-fold. On the one hand, they show the importance of considering active listening behaviour (reaching to sounds and head-movements) in the study of spatial hearing. Methodological approaches based on virtual reality and kinematic tracking, like the one we adopted in the present study, are now more affordable and feasible than in the past. This should motivate researchers to pursue the opportunity to study spatial hearing in more realistic and ecological environments. On the other hand, a better understanding of the mechanisms involved in acoustic space re-learning can have a huge impact on the quality of life of the many hearing-impaired individuals. Specifically, it could pave the way for new approaches for the rehabilitation of spatial hearing difficulties in people suffering from hearing loss or using hearing-enabling devices such as hearing aids or cochlear implants.

## ACKNOWLEDGEMENTS

C.V. was supported by a grant of the Università Italo-Francese (UIF)/Université Franco-Italienne (UFI) and the Ermenegildo Zegna Founder’s Scholarship. F.P. was supported by a grant of the Agence Nationale de la Recherche (ANR-16-CE17-0016, VIRTUALHEARING3D, France) and by a prize of the Fondation Medisite (France). We thank Francesca Guglielmi for helping in collecting data and Giordana Torresani for graphic support (Figure 1A).

